# Vertical growth dynamics of biofilms

**DOI:** 10.1101/2022.08.11.503641

**Authors:** Pablo Bravo, Siu Lung Ng, Kathryn A. MacGillivray, Brian K. Hammer, Peter J. Yunker

## Abstract

During the biofilm life cycle, bacteria attach to a surface then reproduce, forming crowded, growing communities. Many theoretical models of biofilm growth dynamics have been proposed; however, difficulties in measuring biofilm height accurately across relevant time and length scales have prevented testing these models or their biophysical underpinnings empirically. Using white light interferometry, we measure the heights of microbial colonies with nanometer precision from inoculation to their final equilibrium height, producing a novel and detailed empirical characterization of vertical growth dynamics. We show that models relying on logistic growth or nutrient depletion fail to capture biofilm height dynamics on short and long time scales. Our empirical results support a simple model inspired by the fact that biofilms only interact with the environment through their interfaces. This interface model captures the growth dynamics from short to long time scales (10 minutes to 14 days) of diverse microorganisms, including prokaryotes like gram-negative and gram-positive bacteria and eukaryotes like aerobic and anaerobic yeast. This model provides heuristic value, highlighting the biophysical constraints that limit vertical growth as well as establishing a quantitative model for biofilm development.

## Introduction

Biofilms are surface attached microbial communities composed of cells and extracellular polymeric substance (1–3). Biofilms form when a cell attaches to a surface and reproduces, leading to horizontal and vertical expansion. Understanding the dynamics of this expansion, and the processes responsible for them, is fundamental to understanding biofilm development(4–6). Many studies have focused on understanding the horizontal “range expansion,” detailing the impact of physical interactions, the environment, inter-strain competitive dynamics, and more(7–15). Conversely, less is known about the vertical growth of biofilms, despite its importance for determining access to nutrients and oxygen(16, 17). The initial steps of this process (18, 19), as well as some aspects of competitive growth(20–23), have been studied. However, our knowledge of how vertical growth proceeds over short and long time scales is limited. In particular, while growth curves in liquid media have been studied for years building on the works of Jacques Monod(24, 25), and the horizontal expansion of biofilms on agar plates is known to proceed at a constant rate(26–28), we lack detailed measurements and an heuristic understanding of vertical growth. As biofilms play important roles in microbial ecology and human health(29, 30), elucidating this fundamental aspect of microbial physiology is crucial.

The characterization of microbial topographies is a growing field(31–38), but with limited focus on the temporal dynamics of the colony. The lack of clarity about vertical growth dynamics is due in part to the experimental difficulty in measuring the height of a biofilm with sufficient precision over many different time scales in a non-destructive manner(39). Diffraction-limited optical resolution precludes precise measurement of biofilm height at short times. This issue is further confounded by the fact that fluorescent proteins and dyes are specific or localized, and thus do not necessarily report the true height of a biofilm, which is composed of both cells and extracellular matrix. Fluorescent stains can also bleach over time, making it difficult to continuously image the biofilm and the surface it sits on, which is necessary for assessing height. Thus, we lack a clear empirical picture of how vertical growth dynamics proceed over short and long time scales.

Due to the many relevant biological and physical parameters(8, 40–42) involved in biofilm modeling, many models are complex, with multiple interacting phases inside and outside of the colony. Often, these models will incorporate a quadratic carrying capacity term to capture the qualitative features of vertical growth dynamics that are initially rapid before eventually saturating. However, while the logistic growth curve captures these qualitative features, it is unclear if logistic growth is quantitatively accurate. To address this issue, many multi-phase, spatial models have been developed (11, 27, 43–47), with a smaller subset of these models explicitly addressing vertical growth, or colony thickness (9, 48, 49). Even if these models are highly accurate, reliance on many precisely measured parameters limits the practical and heuristic utility of such models. Thus, developing a functional and quantitative understanding of vertical biofilm growth with reduced dimensionality would go a long way towards elucidating this fundamental aspect of microbial physiology.

Here, we measure the vertical dynamics of growing biofilms. Using white light interferometry, we obtain height topographic profiles with single nanometer resolution. We perform frequent measurements for a period of 48 hours, and also test the long term behavior over two weeks. We observe that carrying capacity and nutrient depletion models poorly fit, and are qualitatively inconsistent with, our empirical measurements. Guided by our empirical results, we test a simple interface-limited growth model, which accounts for the fact that diffusion limits the thickness of the growing zone inside the colony. With just three parameters, this model accurately reproduces the observed asymmetric vertical growth dynamics, and captures steady-state biofilm heights at 14 days.

## Results

Using a Zygo Zegage Pro optical profilometer, we measure the growth of multiple microbial colonies, from an initial height of less than a micrometer to large colonies of hundreds of micrometers after 48 hours of growth (Figure 1A-C). These measurements allow us to reconstruct the interface of a growing microbial colony with high resolution across short and long time scales, both in the lateral(0.177 µm) and vertical direction (1 nm) (Figure 1D).

**Figure 1.**
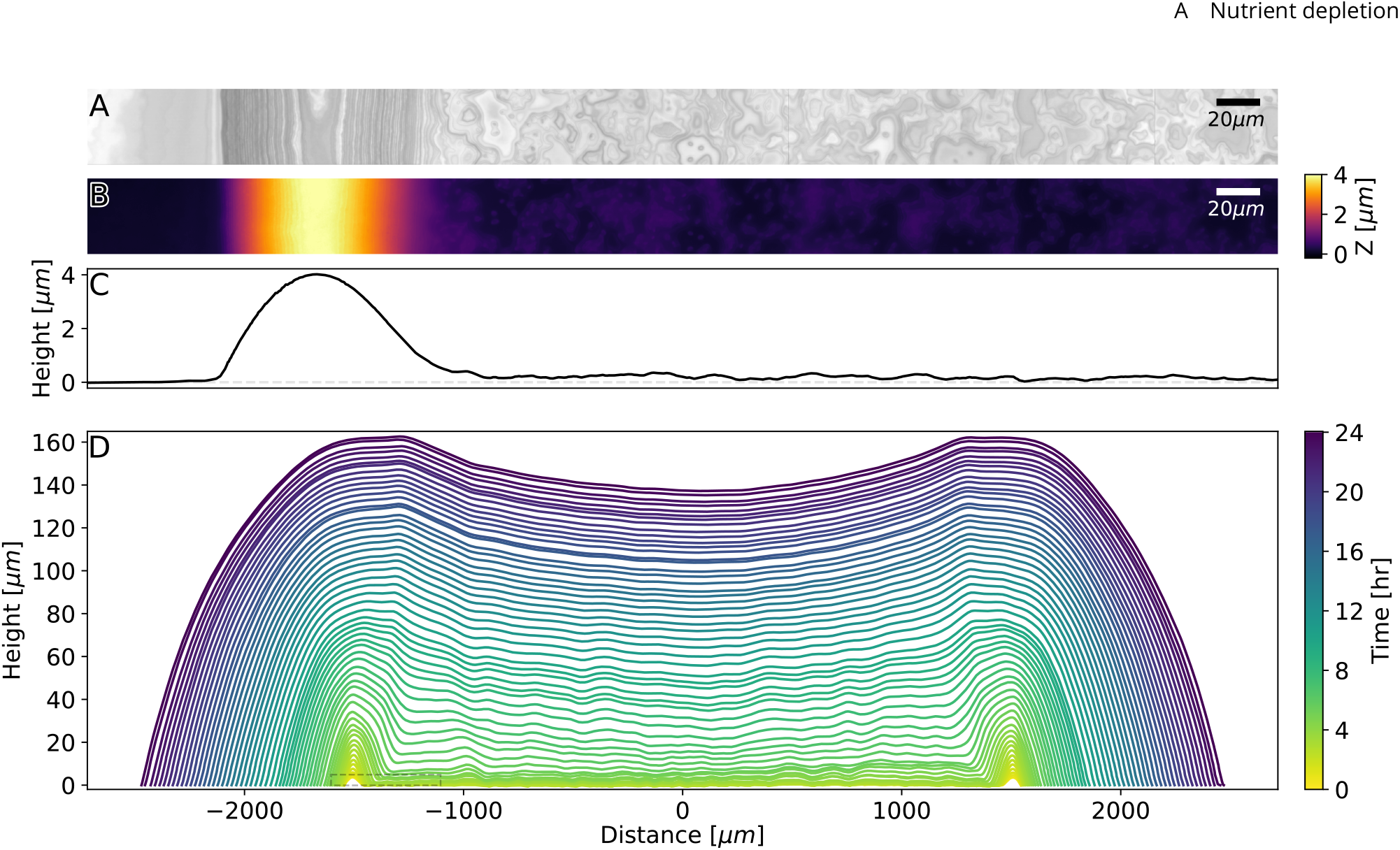
White light interferometry imaging of developing biofilm topography. (A) Light intensity and (B) surface measurement data from the edge of an *Aeromonas veronii* inoculum on LB-Agar are shown 30 minutes after inoculation. (C) One-dimensional averaged profile of the surface topography is computed from the data in (B). (D) A 24-hour timelapse of averaged profiles from growing *A. veronii* biofilm is shown. The colony expands horizontally (x-axis) and vertically (y-axis), with some of its surface features persisting during development. The scale in the y-axis has been increased to better observe the data. The region shown in panels A-C is marked with a rectangle.

We define colony height *h* as the mean height in a 2mm long, 34.6 µm wide strip around the colony center (Figure 1D). Focusing on the center of the colony ensures that we isolate the vertical growth dynamics from the effects of lateral expansion and the initial conditions from inoculation, such as the coffee-ring effect (50, 51). We measured prokaryotic and eukaryotic species, including a wide range of gramnegative and gram-positive bacteria, as well as aerobic and aerotolerant anaerobic yeast (*Saccharomyces cerevisiae*) (see table 2). All species and strains show similar qualitative behavior; we thus initially focus on *A. veronii*, and later expand our study to the full cohort of species and strains.

We observe that the vertical growth rate (i.e., change in height per unit time) changes over time; initially increasing before reaching a maximum and then slowing down, behavior that is qualitatively replicated by logistic growth (Figure 2A). Immediately after inoculation, the vertical growth rate is very low (0-2 µm/hr, see Figure 2A inset). This is consistent with the fact that cells are adapting to a new environment(52, 53) and that doubling events primarily lead to more horizontal colonization until a monolayer is formed(10). During the first few hours of growth, the vertical growth rate increases proportionally with the height (Figure 2B), consistent with exponential growth. The growth rate eventually reaches a maximum value, and then decreases linearly, approaching zero.

**Figure 2.**
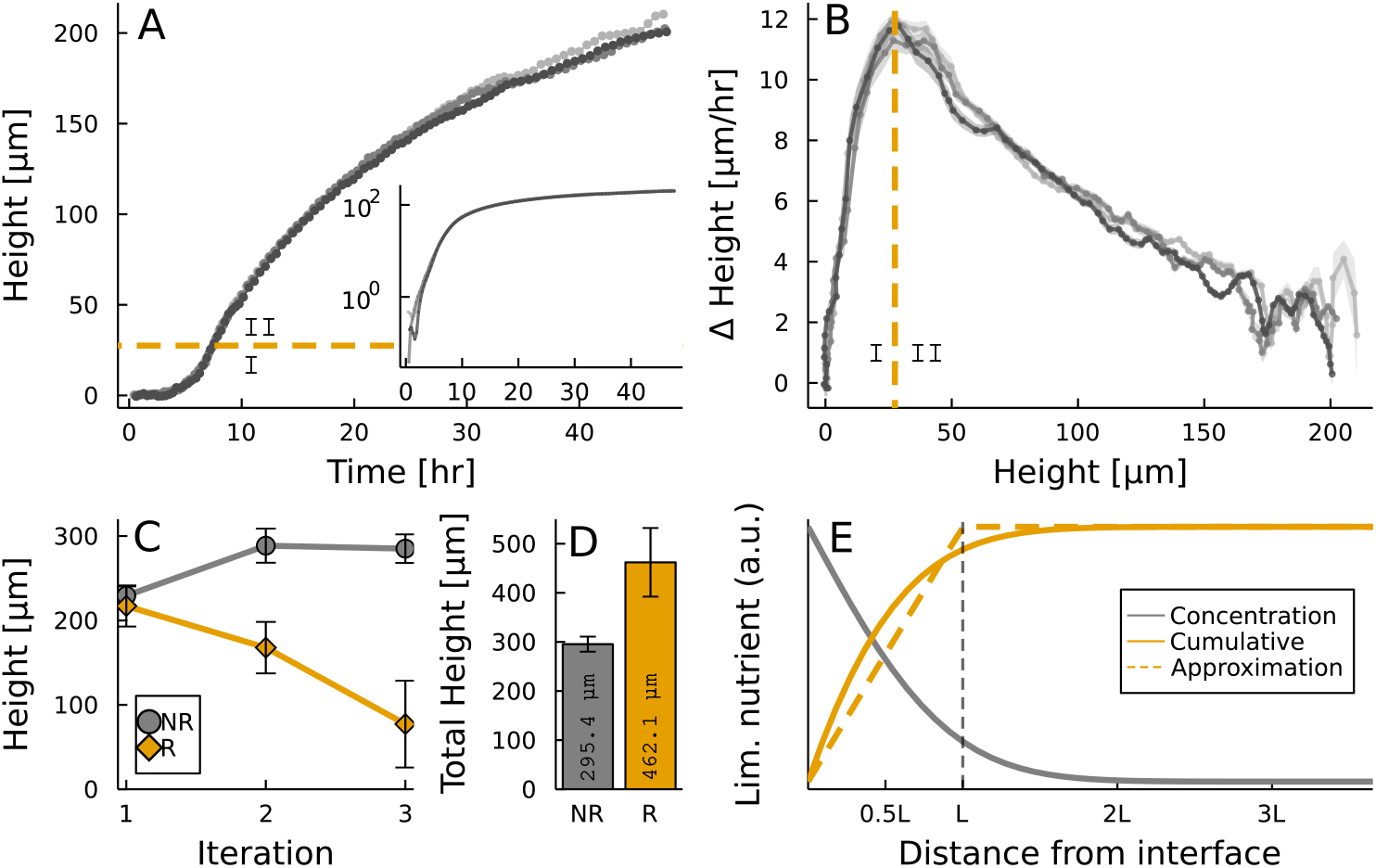
Biofilm height dynamics and geometric constraints. (A) Biofilm height versus time is shown for a 48-hour period in linear and log scales (inset). Three replicates of *A. veronii* are shown. Height increases even at early times, as seen on the log scale. (B) Change in height is plotted against biofilm height. There are two clear regimes: (I) accelerated and (II) decelerated growth, separated by a characteristic length. (C) The height of microbial colonies grown on small agar columns, thus preventing lateral diffusion of nutrients is shown. In one set of experiments, colonies on the agar columns are replaced every two days (R). In a control set of experiments, colonies are not replaced (NR) and instead are allowed to continue growing. (D) The total height of colonies grown on individual agar columns over a period of 6 days is shown. These results demonstrate that colony height does not saturate due to nutreint depletion. (E) Total nutrient concentration through an interface can be approximated as a simple minimum function, and can be used to model the two different growth regimes.

### A. Nutrient depletion

Models of microbial colony growth often directly model the transport and depletion of nutrients. In fact, a prevalent hypothesis is that the biofilm runs out of nutrients, leading to a decrease in growth rate and eventually a final, maximum height. To evaluate the role of nutrient depletion in vertical growth of biofilms, we grew colonies for 48 hours on polycarbonate membranes with pores large enough for nutrients to pass through but small enough to prevent bacteria from passing through, on small agar columns (3.24 cm^2^ in surface area and 0.5cm tall) cut from larger agar plates. These small agar columns limit lateral diffusion of nutrients. On typical large agar plates, nutrients diffuse horizontally and vertically; on these small columns, nutrients can only diffuse vertically up to the colony. After 48 hours, we removed the membrane and colony. We then placed a new membrane on the used column and inoculated, for a second time, in the same location. We found that despite the fact that the first colony nearly reached its saturated height(80%), the second colony grew to 75% of the same height. We then repeated this process on the same plates, i.e., we inoculated at the same location for a third time. Again, we observed that biofilms readily grew, reaching an average height 45% of their previous height. These observations were robust against 12 replicates(Figure 2C). Control plates without replacement showed a total growth of 295.4 µm, while columns with replacement grew a total of 462.1 µm, an excess of 55% (Figure 2D). These experiments demonstrate that vertical growth rates can dramatically slow despite the presence of abundant nutrients. Or, equivalently, in laboratory conditions the steady state height biofilms reach after a long time is not due to the absence of nutrients.

### B. Empirical model selection

We can now use these empirical data to guide the selection of an accurate heuristic model. Plotting Δ*h* as a function of *h* makes it clear that there are two distinct regimes: (I) a linear increase in growth rate with height for *h < L* and (II) a linear decrease in growth rate with height for *h > L* (Figure 2 A-B), where *L* is a characteristic height. The combination of a linear increase and a linear decrease suggests that growth dynamics cannot be governed by a single, unchanging relationship over time. Instead, models of vertical growth must include a “switch” from one regime to another, and a heuristic model of this phenomenon must justify this empirical observation.

One possible explanation for this “switch” from one regime to another comes from nutrient diffusion, the main method of transport inside microbial colonies(54). The key insight is that biofilms only interact with their environment through their interfaces, which limits the nutrient acquisition and posterior diffusion to the cells inside the colony. Nutrients can only enter biofilms through the biofilm-air interface that supplies oxygen to the colony, and the biofilm-substrate interface, which supplies water, carbon sources, and other macroscopic nutrients(55, 56). Ideally, a heuristic model of biofilm growth should be nutrient independent, that is, it should apply equally well to all colonies, regardless of the limiting nutrient. We thus propose a simple “interface growth model” that is biophysically motivated and consistent with empirical data.

To model the impact of these interfaces on nutrient concentration, we directly incorporate the distribution of nutrients as a function of distance from the interface. In fact, the concentration of nutrients near the interface, assuming an unlimited surface source, is known (Figure 2E):

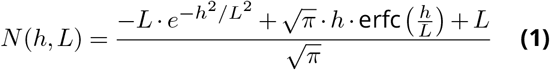

Even with the high resolution of optical interferometry, this level of detail is excessive. First, this analytical expression was derived for infinite media, and biofilms are obviously finite. Second, and more importantly, this relationship does not take into account the active uptake of nutrients. Thus, this analytical expression *N* (*h, L*) does not directly apply to real biofilms.

Nonetheless, equation Eq. S (1) provides us with relevant and critical physical insight. Specifically, the nutrient uptake rate of the biofilms should saturate after the colony reaches a critical height, which we label *h* = *L*. This phenomenon represents the diffusion limit, which prevents biofilms with *h > L* from growing exponentially. To incorporate these biophysical insights and account for the uncertainty in the exact functional form of *N* (*h, L*) for real biofilms, we approximate this relationship as min(*h, L*), a simple linear function that reaches its maximum value of *L* for *h ≥ L*. Similar expressions have been previously derived and explored in mathematical models for biofilm growth (9, 57). This simplified expression captures, to the first order in *h*, the underlying biophysics of diffusion.

We now assemble a heuristic model of vertical growth that takes these empirical and biophysical phenomena into account. These phenomena can be captured with a simple 3-parameter differential equation:

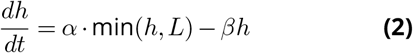

Where *α* is the growth rate, *β* is the decay rate, and *L* is the diffusion length of the limiting nutrient. This model can be rewritten as a piecewise function in relation to the critical height *L*:

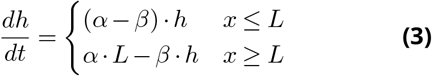

This expression captures the main biophysical processes governing vertical growth, while remaining simple for any implementation or quick back of the envelope calculations.

### C. Model accuracy

We now test the accuracy of the interface limited model, as well as two minimal models that rely on common alternative approaches to modeling vertical growth: logistic and nutrient depletion models. These reference models do not represent the state of the art, but instead provide a simple implementation of common modeling choices, allowing us to test their accuracy. Specifically, the logistic model is built on the commonly used carrying capacity term that is quadratic in *h*, and the nutrient depletion model incorporates first-order growth and death terms, with a nutrient field that is consumed over time. We will also consider logistic and interface models that incorporate nutrient fields, to test if doing so results in improved accuracy (for more details on these alternative models, see Materials and Methods).

Over 48 hours of growth at room temperature, the interface model accurately captures colony height across the full time period (Figure 3A). This is true both when the auxiliary nutrient field *c* is or is not included, supporting the idea that nutrient depletion is not the limiting factor for growth, and does not need to be directly incorporated for model accuracy. The root-mean-squared error, RMSE, quantifies the relative agreement of these models: 0.15 µm for the interface model without a nutrient field, and 2.71 µm for the interface model with a nutrient field (Figure 3A inset). This good agreement can also be quantified with the residual as a function of time (Figure 3B). The interface-limited model oscillates around 0 µm, with fluctuations similar to the expected differences from the 3 parallel replicates. Further, the alternative models are poor fits for the height dynamics; these models grow slower than the experimental data at early times, and then reach their saturated height too early (Figure 3A). For each of the alternative models RMSE>10 µm.

**Figure 3.**
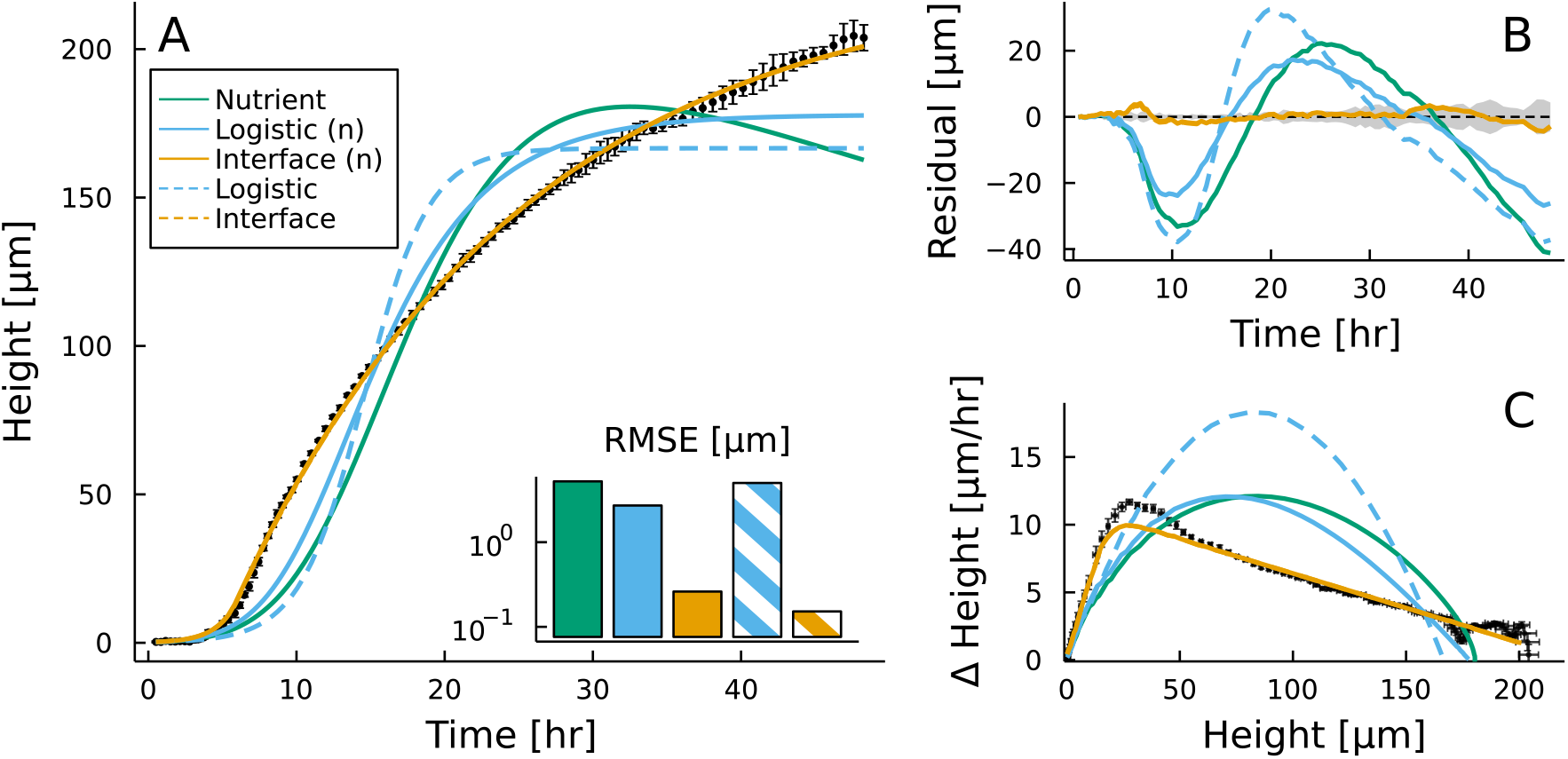
Interface model outperforms the nutrient and logistic models over 48 hours of growth. (A) The mean height of *A. veronii* colonies, averaged over three replicates, is shown versus time. Error bars represent standard deviation across replicates. Best fit lines for the interface, logistic, and nutrient models are shown. Interface and logistic models expanded to include depletable nutrient fields are shown in solid lines; models without nutrient fields are shown as dashed lines. Note, the two interface models overlap in the figure, but the model values do differ across the data, as can be seen by their different RMSEs. (B) Residuals for the best fit predictions from the above models are shown as a function of time. On average, the nutrient model is off by 15.47*µm*, the logistic model by 10.77*µm*, and the interface model by 1.24*µm*. The gray region corresponds to the standard deviation of the 3 replicates in relation to the mean value at each time. (C) Change in height, Δ Height, is shown as a function of Height for best fit predictions. The two interface models follow the empirical data much more closely than all other models.

To determine why the alternative models fail, we examine the change in height as a function of the current height for all models (Figure 3C). The interface limited model directly incorporates the empirical relationship between Δ*h* and *h*, and thus accurately fits these data. Conversely, the alternative models completely fail to capture the observed linear relationships between growth rate and height.

### D. Behavior on long time scales

We next test the agreement of the interface model on long time scales, measuring *h* over a period of two weeks. To do so, we measure the growth of three different species representing a wide range of microbial growth dynamics: (i) the previously introduced *A. veronii*, (ii) *E. coli*, and (iii) an aerotolerant anaerobic *S. cerevisiae* mutant. The interface model accurately fits the first 48 hours of vertical growth for *E. coli* and the anaerobic *S. cerevisiae*, with RMSE of 1.7 µm and 3.07 µm, respectively. To test long term accuracy, we inoculate multiple colonies on multiple plates, and then measure all colonies on a new plate every two days (see Materials and Methods for more details). The interface model accurately captures the vertical growth dynamics of all three microbes on long time scales (Figure 4A), while exhibiting a linear decrease in growth speed, consistent with the interface model, although at different heights and time scales (Figure 4B).

**Figure 4.**
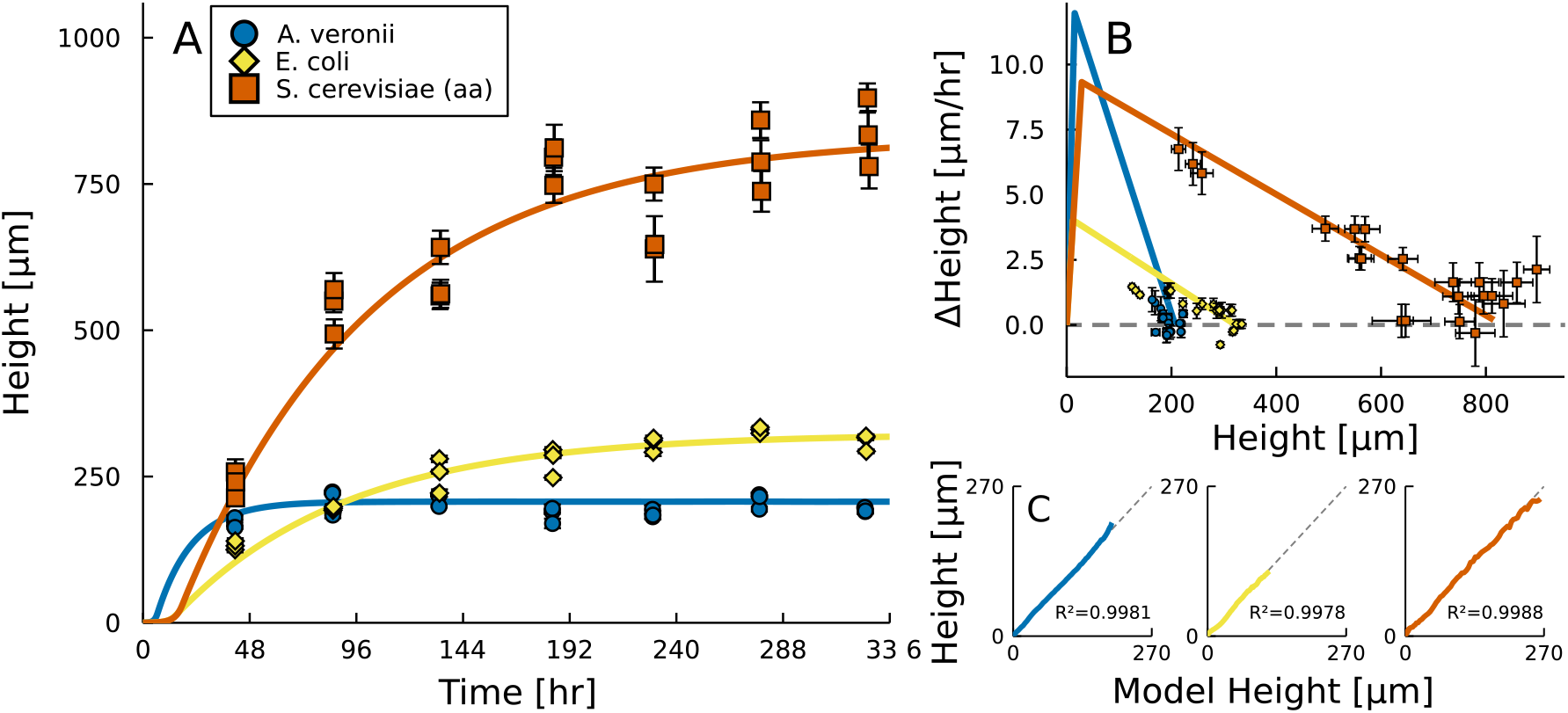
Long term growth of microbial colonies. (A) Measurements of height versus time are shown for three different species. Error bars represent standard deviations across the 2mm homeland region. Solid lines show the best-fit interface model. (B) Change in height is shown versus colony height in the 2-14 day range. The slow linear decrease in growth rate is observed in all measured strains. The solid line represents the best-fit growth rates according to the interface model. (C) Colony heights during the initial 48 hours of growth of plotted against best fit predictions from data taken during the 2-14 day range. The agreement between the data and the model is evident.

It is important to note that the long time agreement of the interface model does not come at the cost of accuracy on short time scales (Figure 4C). We compared the experimental data and the model predictions in the 48-hour period; for all three colonies *R*^2^ was above 0.99. The ability to capture short and long term dynamics, simultaneously highlights that, in fact, vertical growth is accurately described by the simple interface model.

We next test the predictive power of the model at short times to determine what governs the dynamics and steady state behavior of long-standing colonies. These three species exhibit different growth rates, and different heights after the initial 48 hours of growth: 203.83 ± 4.34, 208.64 ± 8.07, and 116.88 ± 5.33 µm for *A. veronii, E. coli*, and aerotolerant anaerobic *S. cerevisiae*, respectively. The predicted steady state maximum height *h*_max_ can be obtained from the model parameters, following the expression:

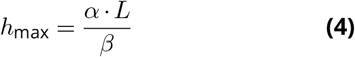

This expression correctly predicts that *S. cerevisiae*, will continue to be the tallest colony by a considerable amount, and, more interestingly, that *E. coli* will surpass *A. veronii* in height, despite the fact that *A. veronii* is almost twice as tall as *E. coli* at 48 hours. This prediction would not be possible without high resolution data or the accurate model.

However, while experimental steady state heights fall in the 95% confidence interval of the interface model fitted using the initial 48 hours of data(Figure S7), the range of this interval varies substantially across these three species (21.6 µm, 215.4 µm, and 473.23 µm for *A. veronii, E. coli*, and aerotolerant anaerobic *S. cerevisiae*, respectively).

The source of this wide range of uncertainties is the difficulty in rapidly measuring *β. β* is small, on the order of 10 nm/hr. 48 hours is thus not necessarily enough time for an accurate measurement, leading to a wide range of effective growth rates at the end of the measurement. A colony that is already close to its steady state height (*A. veronii*) has a much narrower confidence interval than a colony that is still growing rapidly (*S. cerevisiae*). To determine *β* with narrow confidence intervals, a large amount of data in the *h > L* regime is necessary.

To demonstrate this issue, we performed an 88 hour long timelapse of *E. coli*, and obtained the best fit parameters as a function of time. It is clear that by 40 hours *α* has already converged (Figure S8), whereas *β* and *L* do not converge to their final values until ∼ 65 hours. It is important to note that the percentage change of these values varies greatly, with the limiting parameter being *β*, which at 50 hours is 230% of its final steady state value. On its own, fluctuations in *β* that range from 10-23 nm/hr do not seem substantial. However, in light of Eq. S (4), these fluctuations can greatly change the predicted final height of the colony. As a result, *h*_max_ cannot be accurately predicted until *β* is accurately measured.

### E. Universality of the vertical growth dynamics of the interface growth model

Finally, while we have thus far focused on a small number of species, the interface growth model is built on biophysical limits of diffusion that should readily apply to many species and strains. To test how broadly this model applies, we perform measurements on nine different organisms including (1) different cell sizes (from ∼ 1 - 10 µm), shapes (from rods to commas to nearly spherical ellipsoids), (3) different extracellular matrix production (from engineered strains with limited extracellular matrix production to wild type *V. cholerae* that are known biofilm formers), (4) prokaryotic and eukaryotic species, (5) gram positive and gram negative species, and (6) aerobic and anaerobic fungi.

For this purpose, we measure the vertical growth of these microbes, with three parallel replicates, for a period of 48 hours. This length of time is sufficient to capture the two growth regimes delimited by *L*, and thus fit all interface model parameters. We observe excellent agreement between the interface model and the vertical growth dynamics of all investigated species and strains (see Figure 5), despite dramatic differences in colony morphologies and growth rates during these 48 hours of measurement.

**Figure 5.**
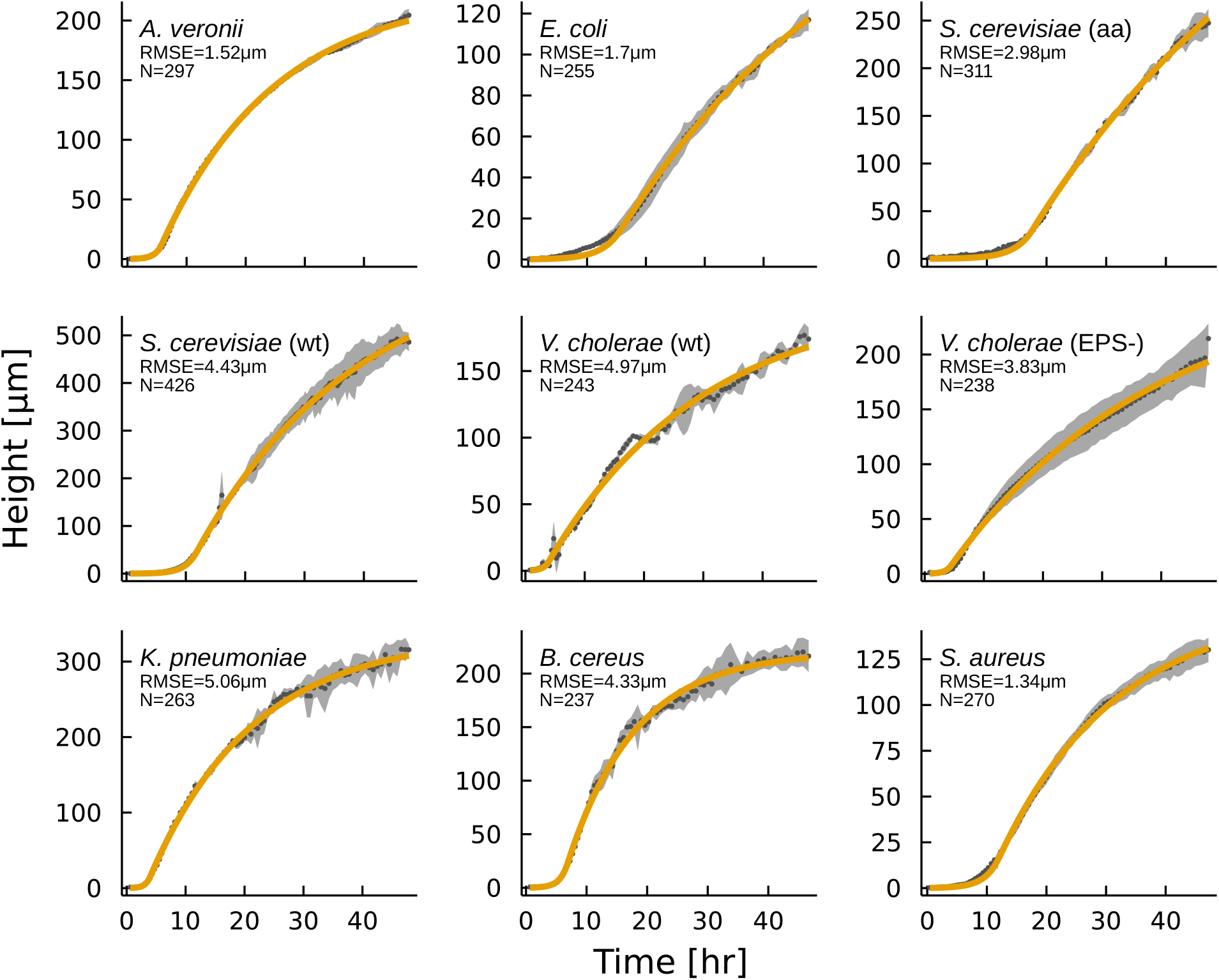
Growth of different species and strains over 48 hours. The average height plus or minus the standard deviation across 3 parallel colonies is shown in gray. Error bars represent standard deviation across replicates. The best fit interface model is shown in orange. The model RMSE and the total number of interferometry profiles analyzed are reported in each panel. While these microbial colonies differ in their composition, height, radius, and overall morphologies, the interface model accurately describes the average height dynamics over time for each one.

We also summarize the allowed ranges for growth parameters in table 1. This resource is useful for future modeling work, with tested and empirical evidence on the accuracy of the model. The parameter range across the sampled cohort is quite large. *α* ranges by a factor of 4, between 0.3 hr^-1^(*E. coli*) to 1.3 hr^-1^ (*K. pneumoniae*). *L* also exhibits a wide range of values, from 8 µm (both *V. cholerae* strains) to 45 µm (*S. cerevisiae)* grown on YPD media). *β* is more difficult to characterize. Its range of best fit values is consistent across different strains, going from 0.02 hr^-1^ to 0.05 hr^-1^ for *E. coli* and *B. cereus*, respectively. However, the confidence intervals, relative to the best fit parameter value, are typically larger for *β* than for *α* and *L*. Due to their large sizes, the true *β* values may either be nearly all ∼ 0.04 hr^-1^, or distributed across a range from 0.01 hr^-1^ to 0.07 hr^-1^.

**Table 1.** Interface model parameters for the cohort of microbes. Shown is the best-fit, and 90% Confidence interval for the 48 hour data.

| Strain | $\alpha$[1/hr] | $\beta$[1/hr] | $L$[μm] |
| --- | --- | --- | --- |
| *Aeromonas veronii* | 0.875(0.858 – 0.911) | 0.051(0.048 – 0.055) | 13.205(12.277 – 13.878) |
| *Escherichia coli* | 0.339(0.326 – 0.360) | 0.023(0.012 – 0.036) | 14.347(11.705 – 17.622) |
| *Saccharomyces cerevisiae* (aa) | 0.345(0.337 – 0.350) | 0.023(0.011 – 0.028) | 32.071(27.485 – 33.612) |
| *Saccharomyces cerevisiae* | 0.555(0.538 – 0.579) | 0.037(0.026 – 0.05) | 44.018(37.053 – 51.257) |
| *Vibrio cholerae* (wt) | 1.112(0.99 – 1.284) | 0.038(0.031 – 0.048) | 7.066(5.559 – 8.861) |
| *Vibrio cholerae* (EPS-) | 0.995(0.886 – 1.232) | 0.029(0.017 – 0.045) | 7.783(5.371 – 10.532) |
| *Klebsiella pneumoniae* | 1.316(1.253 – 1.440) | 0.057(0.05 – 0.064) | 14.475(12.241 – 16.278) |
| *Bacillus cereus* | 0.881(0.858 – 0.921) | 0.089(0.041 – 0.107) | 22.27(18.486 – 25.641) |
| *Staphylococcus aureus* | 0.475(0.464 – 0.488) | 0.053(0.041 – 0.063) | 17.025(14.726 – 18.783) |

## Discussion

We investigated the vertical growth dynamics of microbial colonies using timelapse interferometry. This technique allows continuous non-disruptive measurements across broad time and length scales with single nanometer resolution, enabling us to characterize the vertical growth dynamics of microbial colonies with unprecedented precision. Using these high precision data, we found that the vertical growth rate initially increases linearly with height; after reaching a critical length scale (*L*), the growth rate then decreases linearly with height. We show that a simple heuristic model built on the interface-limited diffusion of nutrients accurately captures short and long term dynamics. This minimal model was validated across many different species and strains of bacteria and fungi.

The use of interferometry was essential for the empirical characterization of vertical growth dynamics presented here. For example, high resolution measurements were necessary to identify that growth rate decreases linearly with *h* for *h > L*, rather than quadratically with *h*. This empirical relationship provides valuable insight into the development of microbial colonies. Further, *β*, the linear decay rate, is typically very small, on the order of tens of nanometers per hour. These measurements were possible as interferometry readily provides both the high resolution out-of-plane necessary to capture small changes in height, as well as the fast measurement speed necessary to characterize changes over time.

Interferometry, however, has three key limitations. First, this technique only measures the top surface of a colony, and cannot measure the internal structure. Second, the steepest measureable slope is limited by the numerical aperture of the objective used. When the surface is too steep (*θ* > 28.13° for the objective we used) the light path does not return to the objective. This slope threshold may limit the use of interferometry on colonies that are highly wrinkled or buckled(58–60). Finally, optical interferometry requires that a surface be sufficiently reflective. This condition will always be met for the colony-air interface; however, measurements at biofilm-fluid interfaces, while possible, are much more difficult. Despite these limitations, interferometry measurements work in a broad range of surface attached colonies and allow different models of vertical growth dynamic to be tested.

Models of biofilm growth are diverse; they incorporate different biophysical phenomena and use different mathematical approaches. However, the lack of empirical data characterizing vertical growth dynamics prevented these models and their underlying assumptions from being tested. Crucially, the empirical data presented here have the resolution necessary to test the accuracy of existing models, evaluate their underlying assumptions, and then use empirical data to develop an accurate heuristic model.

For example, we find that models built on logistic growth fail to capture the correct relationship between growth rate and colony height. A logistic term is functionally equivalent to a term that decreases growth rate quadratically with height. This term is mathematically appealing; it is the lowest order term that qualitatively describes the changes in growth. Despite its conceptual appeal, it lacks empirical justification (Figure 3C). Of course, most of the models developed in the literature are complex, and include more terms and fields in the set of governing equations. However, the evidence presented here suggests that going forward, logistic-like vertical growth should be replaced with empirically-supported expressions.

Similar to logistic growth, we found that nutrient depletion models are not empirically supported, since they do not capture the relationship between growth rate and height(Figure 3C). Further, for rich media laboratory experiments, nutrient depletion is not responsible for the eventually cessation of vertical growth (Figure 2 C-D), and adding nutrient depletion to the interface model did not improve their accuracy(Figure 3A). While nutrient depletion may be important to account for near-starvation conditions, it is not a critical component of vertical growth in typical laboratory conditions.

Empirical measurements of growth rate as a function of height demonstrate that three terms are necessary to fully describe vertical growth dynamics. The inclusion of each of these three terms in the interface model we propose is motivated by three fundamental biophysical quantities. *L* is the length at which growth rate ceases to increase in proportion to biofilm height. This term is inspired by, and consistent with, the length at which diffusion limits the transportation of nutrients through biofilm interfaces. The growth rate *α* captures all vertical growth, including the growth and doubling of cells, but also secretion of extracellular substances. The decay rate *β* captures all effects that decrease growth rate, such as mechanical settling, cell death and lysis, cells entering a lag phase, and diminished efficiency in the reabsorption of biomass. In other words, *α* and *β* coarse grain all phenomena that lead to an increase or decrease (respectively) in growth rate into two simple terms. Rather than limiting the utility of this model, coarse graining places the emphasis on the net impact of increased height on growth rates, which may be universal, rather than on the microscopic mechanisms that modify growth rates, which will often be specific and heterogeneous. This approach provides a useful heuristic model for vertical growth dynamics, as opposed to more complex alternatives(61).

The unique insight provided by high resolution studies of vertical growth dynamics is exemplified through two case studies comparing: (i) the role of respiration in *S. cerevisiae*, and (ii) the role of extracellular matrix production in *Vibrio cholerae*. First, we find that there is no qualitative difference between the vertical growth dynamics of aerobic and obligate anaerobic strains of *S. cerevisiae*. The aerobic yeast strain used in this study is “mixotrophic,” capable of both fermentation and aerobic respiration(62) depending on the concentration and types of sugars found in the environment. The anaerobic strain has lost part of its mitochondrial genome and is therefore only capable of fermentation, which is not oxygen-dependent. Multicellular anaerobic yeast have recently been shown to overcome group size constraints that aerobic yeast face due to the oxygen diffusion limitation into the center of a group(63). The aerobic strain grows faster than the anaerobic strain (63), but in each case the vertical growth dynamics proceed following the interface model. In a different vein, we directly compare the vertical growth dynamics of strains of *V. cholerae* with and without EPS. Vibrio polysaccharide is an essential component for biofilm formation, and the associated genes are controlled by the master biofilm regulator, *VpsR*(64, 65). Deletion of *vpsR* (EPS-) prevents biofilm production in *V. cholerae*, leading instead to the growth of “smooth” colonies with much less extracellular matrix than wild type strains. Again we find that the vertical growth dynamics do not change substantially in the presence or absence of EPS. Thus, microbial colony vertical growth dynamics appear relatively universal, even across microbial strains with different compositions.

As the interface model is a simple heuristic model for vertical growth, there are many details it does not address. For example, the model predicts the dynamics of the bulk height, and does not distinguish between cells and extracellular matrix within the colony. Relatedly, as the model focuses on the mean total height it does not address spatial fluctuations. Topographic fluctuations likely encode insight into the underlying phenomena behind vertical growth dynamics, as previously demonstrated in homeostatic biofilms (66, 67). In a different vein, the model does not account for more complex environmental conditions, such as limited access to nutrients(68), environmental stresses(69), or heterogeneities in the surface on which the colony grows(70). Further, in nature biofilms often contain multiple interacting species and subpopulations(71), leading to complex dynamics and structuring(5, 72) of the colony that we do not consider. The work presented here provides a foundation on which these detailed questions about vertical growth dynamics can be studied.

The study of microbial community interfaces via interferometry presents an easy and inexpensive approach to studying the structure of microbial colonies. Interfaces and surface fluctuations have long been studied in physics, both for the information they provide about what occurs beneath the surface as well as the rich phenomena that surfaces themselves exhibit. Indeed, biology also has a rich history of studying interfaces, from the surfaces of cells or organs(73–76), to biome-scale interfaces(77). Biological interfaces on meso-scales are traditionally less studied, despite their known importance in a wide range of systems(78–80) including bacterial communities (81, 82). This is likely due to the difficulty in measuring these microscopic, and often sticky, interfaces. Other techniques for characterizing the height and topographies of colonies lack the resolution (e.g., confocal microscopy), are slow and potentially destructive (e.g., atomic force microscopy), or do not readily permit time lapse measurements (e.g., scanning electron microscopy)(39). While these alternative measurement techniques might be modified to overcome their limitations(83), interferometry is already very well-suited for studying biofilm interfaces, as demonstrated here and in previous studies (32, 66, 67). These works represent the first attempts in utilizing optical interferometry as a tool for microbiology, an approach that will only get more useful when paired with the development of heuristic models and detailed measurement protocols.

Finally, the dynamics of vertical growth represent a fundamental aspect of microbial physiology. Biofilms are exceedingly common; they are found in many environments (84–86) and are of key importance from a medical(87–91) and economic(92) perspective. An understanding of how biofilms develop their 3-dimensional structure is thus relevant for diverse questions of microbial ecology, evolution, and human health. In particular, understanding vertical growth dynamics is crucial for assessing the underlying biophysical process regarding biofilm development: the way resources do and do not limit growth, the relative fitness of various biofilm morphologies (e.g., thin biofilms vs. thick biofilms). The vertical growth measurements and model describing them are but one of the many aspects in biofilm development, but necessary ones for proper understanding how these microbial colonies grow.

## Methods and Materials

All inoculations consist of 1.5 µL from an overnight culture grown at 37 °C diluted to *OD*_600_ = 1. Control plates were left to grow at a temperature of 23.8 °C. For interferometry measurements, we designed an enclosure that allows continuous measurements with minimal media and sample evaporation. The full enclosure maintains a temperature of 23.8 °C and a relative humidity of 80% for the full measuring period.

Since colony growth is radially symmetric, we only measure a horizontal strip of the biofilm, allowing us to achieve better temporal resolution. The measurement proceeds across the entire colony, and includes uncolonized substrate surface on both sides of the colony. Given the aspect ratio of our data and the goals of our study, we then lower the dimension of the data by averaging on the Y-axis (Figure 1A-C). Then, we utilize the substrate height measurements to obtain the relative height of the colony to its background.

### Column growth

We label and cut columns of LB-agar to dimensions of 18mm× 18mm × 5mm. On each column we deposited a Polycarbonate Track Etch Membrane with 0.2 µm pore size, 13mm diameter (GVS brand). We inoculate 1.5µL of OD1 bacterial suspension on top of said membrane. This is repeated over 6 plates (18 columns/colonies). Every 48 hours, up to 3 iterations, we measure the growth on each column. On 9 of the columns we than removed the membrane and deposited a new membrane and inoculated on top of the new membrane.

### 48h timelapses

We inoculated 3 colonies of the measured strain, on two separate petri dishes with LBAgar. One plate was measured continuously (replicates A-C) for 48 hours in the profilometer enclosure. Measurements consist of horizontal radial strips of each colony with a Zygo ZeGage Pro optical profilometer with a 50x Mirau objective (NA 0.55), on the 1000×200 @800Hz capture mode. We increase the Field of View of the measurement adaptively, to always capture an uncolonized region outside the growing colony. Colonies that were continuously measured did not show any significant growth differences compared to their sealed counterparts (replicates D-) after the experiment (FIGSUPPinterferometrycontrols).

### Long time measurements

We inoculated 21 plates 7 for ZOR0001, 7 MG1655, and 7 GOB33 (table 2), with 3 separate colonies each. All plates were sealed with parafilm after inoculation. Every two days, we performed radial measurements with a 10x Mirau objective (NA 0.3) 1000×200 @800Hz capture mode. Each plate was discarded after measurement.

**Table 2.** Timelapses overview details.

| Strain | Species | Media | Details | Cell shape | Gram |
| --- | --- | --- | --- | --- | --- |
| ZOR0001 | *Aeromonas veronii* | LB 1.5% | wild type | rod | negative |
| MG1655 | *Escherichia coli* | LB 1.5% | wild type | rod | negative |
| GOB33 | *Saccharomyces cerevisiae* | YPD | petite yeast | ellipsoid | - |
| Y55 | *Saccharomyces cerevisiae* | YPD | wild type | ellipsoid | - |
| C6706 | *Vibrio cholerae* | LB 1.5% | wild type | comma | negative |
| C6706 | *Vibrio cholerae* | LB 1.5% | $\Delta vpsR$ | comma | negative |
| Top52 | *Kelbsiella pneumoniae* | LB 1.5% | wild type | rod | negative |
| SW520 | *Bacillus cereus* | LB 1.5% | wild type | rod | positive |
| SW519 | *Staphylococcus aureus* | LB 1.5% | wild type | spherical | positive |

**Table 3.** Fitting boundaries for model fitting. Ranges push the boundaries of what makes physical/biological sense by at least an order of magnitude.

| Model | $\alpha[1/hr]$ | $\beta[1/hr]$ | $L[\mu m]$ | $K[\mu m]$ | $K_c[a.u.]$ | $\epsilon$ |
| --- | --- | --- | --- | --- | --- | --- |
| Nutrient (n) | $[10^{-5}, 10^3]$ | $[10^{-5}, 10^2]$ | - | - | $[10^{-5}, 10^3]$ | $[10^{-5}, 10^5]$ |
| Logistic (n) | $[10^{-5}, 10^3]$ | - | - | $[1, 10^5]$ | $[10^{-5}, 10^3]$ | $[10^{-5}, 10^5]$ |
| Interface (n) | $[10^{-5}, 10^3]$ | $[10^{-5}, 10^2]$ | $[1, 10^5]$ | - | $[10^{-5}, 10^3]$ | $[10^{-5}, 10^5]$ |
| Logistic | $[10^{-5}, 10^3]$ | - | - | $[1, 10^5]$ | - | - |
| Interface | $[10^{-5}, 10^3]$ | $[10^{-5}, 10^2]$ | $[1, 10^5]$ | - | - | - |

#### Geometric constraint derivation

The geometric constraint detailed in Eq. S (1) is rooted in a simple diffusion equation. This expression is used in fabrication principles and was taken from (93). Starting from Fick’s Law for a concentration field *c* and diffusion coefficient *D*:

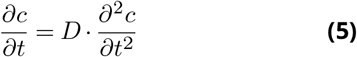

By direct integration we obtain that the distribution of nutrients as follows:

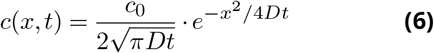

where *c*_0_ is the initial concentration. If we now consider an infinite source with concentration *c* = *c*_0_ on one side of a slab of material (in our case, a microbial colony), the concentration on a finite element of width *dξ* is:

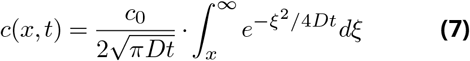

Using the error function

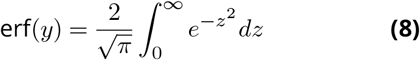

we can rewrite the expression as:

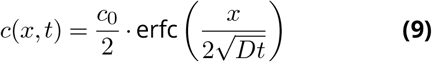

where erfc is the complimentary error function. Finally, we can obtain the total amount of nutrients in a finite sized slab of height *h* by integrating this expression:

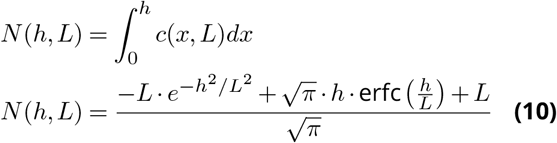

where we removed the temporal dependence by a characteristic diffusion length 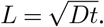.

#### Growth models and ODE simulations

As benchmarks for comparison to the interface model we utilize two intuitive models, a nutrient depletion model and a logistic model.

### Nutrient depletion model

This model captures the dynamics of how a biomass of an available nutrient *c* is transformed into biomass height *h* (assuming that mass∝height in a column). Both fields are linearly related; therefore a decrease in growth rate has to be due to the lack of nutrient availability.

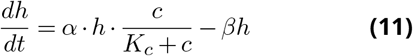

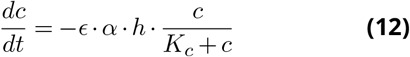

Here, *α* is the growth rate, *β* the decay rate, 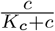 is the Monod kinematic term for nutrient uptake, and *ϵ* is the inverse conversion rate for the transfer. For our simulations we set *h*_0_ = 0.01*µm*, and *c*_0_ = 1.0. This model can not be used without nutrient *c* dependency, since the equation would then be reduced to the monotonic equation 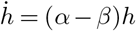

### Logistic

The use of a logistic model makes sense in a qualitative manner, since we do observe a behavior that follows “accelerated-decelerated” growth. The issue with these kind of models is that while simple, they associate the negative(−) term in the differential expression to a quadratic term, obscuring the meaning of the variable.

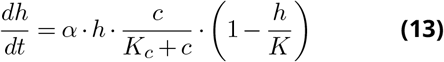

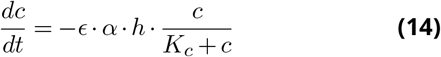

Variables are the same as in the previous model, with *K* representing the carrying capacity, or the maximum height the colony can reach. We can also utilize the logistic model without a nutrient dependency, and rewrite the expression as 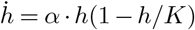.

#### Parameter estimation

We optimized the multiple differential equations against experimental data with a sum of squared error loss *J*:

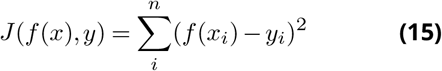

where *y* is the collection of experimental heights corresponding to measurement times *t, f* (*x*) is the evaluation of the differential equation at time *t*. This allow us to directly compare the simulated predictions with our non equidistant temporal measurements. While this loss function *J* prioritizes better agreement with the data on higher heights, results do not suggest that compensating with higher weights at early times is needed since residual values are low across all the measurement period.

To get confidence intervals and parameter distributions we utilize a moving block bootstrap across the 3 different replicates, sampling n=20 blocks of size s=5, returning a total of 100 points, roughly the same amount as an individual timelapse. We iterate for i=1:1000 times and obtain the best-fit parameters *α*_*i*_, *β*_*i*_, *L*_*i*_. Then utilizing equation Eq. (4), discard the outlying 5% values.

All parameter estimation was done in Julia with the DifferentialEquations.jl(94) and DiffEqFlux.jl(95) packages.

## Data availability

Source code for the cleaning, analysis and figures, and clean datasets have been uploaded to a [Github repository](https://github.com/pbravoc/vertical_growth_biofilms). Due to file size limits, full unprocessed images are available upon request.

## Acknowledgments

P.J.Y. acknowledges funding from the NIH NIGMS (Grant No. 1R35GM138354-01) and NSF Biomaterials (Grant No. BMAT2003721). B.K.H. acknowledges funding from NSF Biomaterials (Grant No. BMAT2003721).

## Supplementary Information

**Figure S1.**
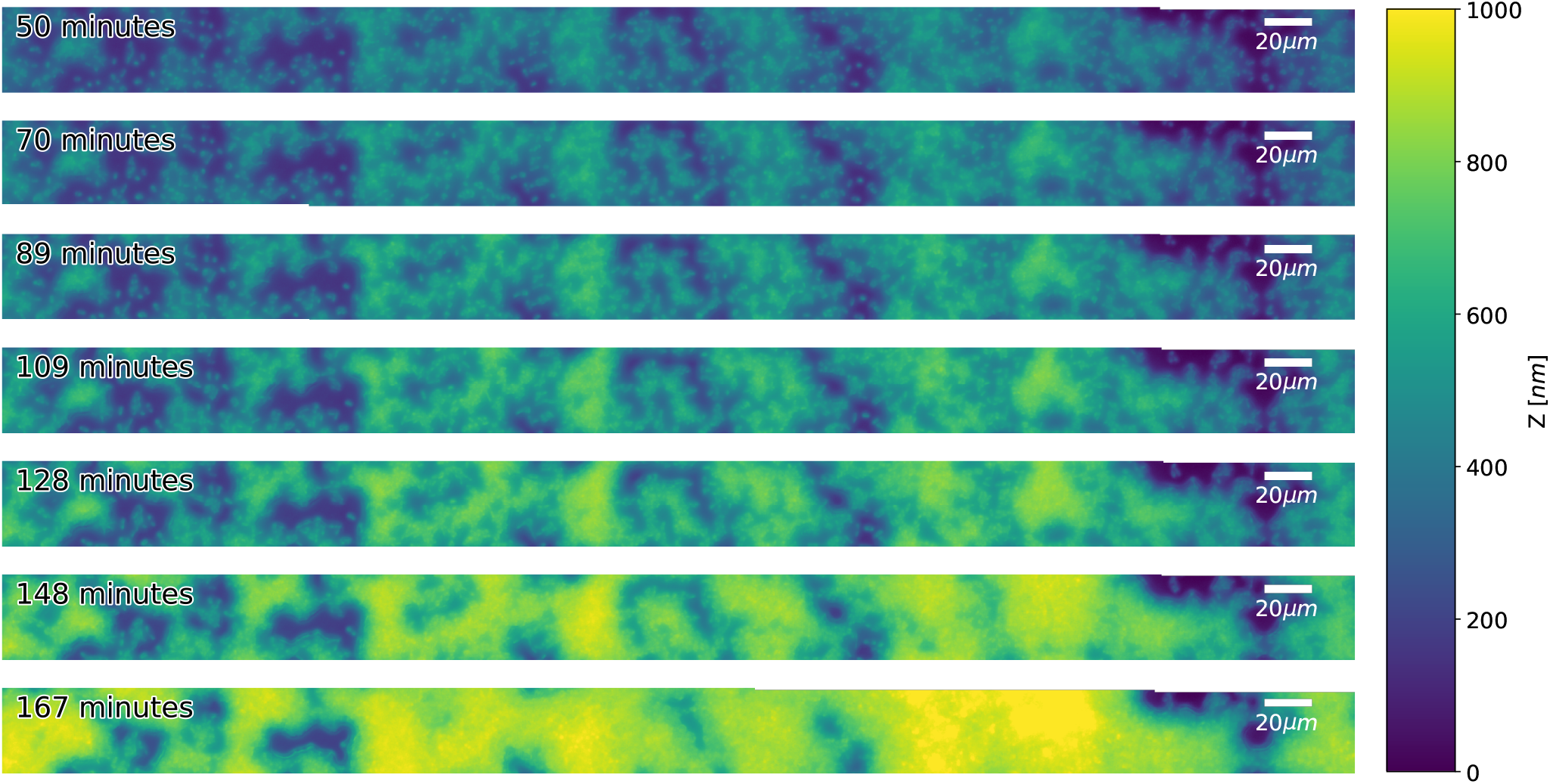
Interferometry allows accurate measurement of the spatial vertical growth dynamics. An *A. veronii* colony after inoculation. Initially, individual cells can be observed undergoing reproduction through their height profiles.

**Figure S2.**
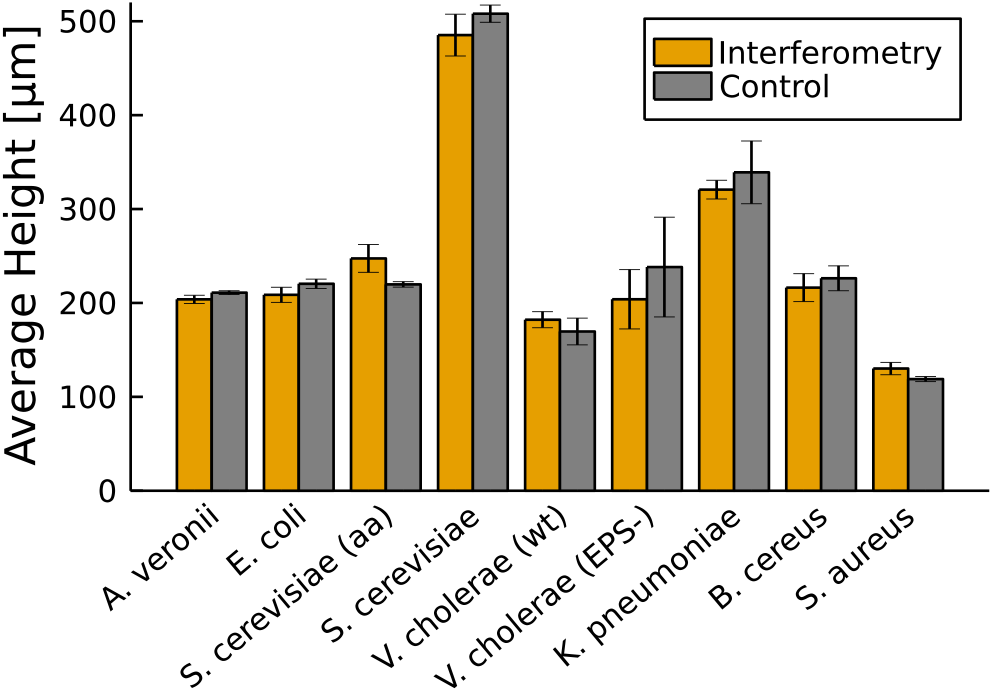
Colony heights are not affected by continuous measurement under the interferometer. Control samples were inoculated from the same overnight batch on a different plate, which was then left to grow at room temperature, sealed with Parafilm. Comparisons were after each timelapse finished, at 48.6, 87.7, 47.6, 48.6, 49.5, 50.2, 48.3, 47.1, 47.48 hours respectively.

**Figure S3.**
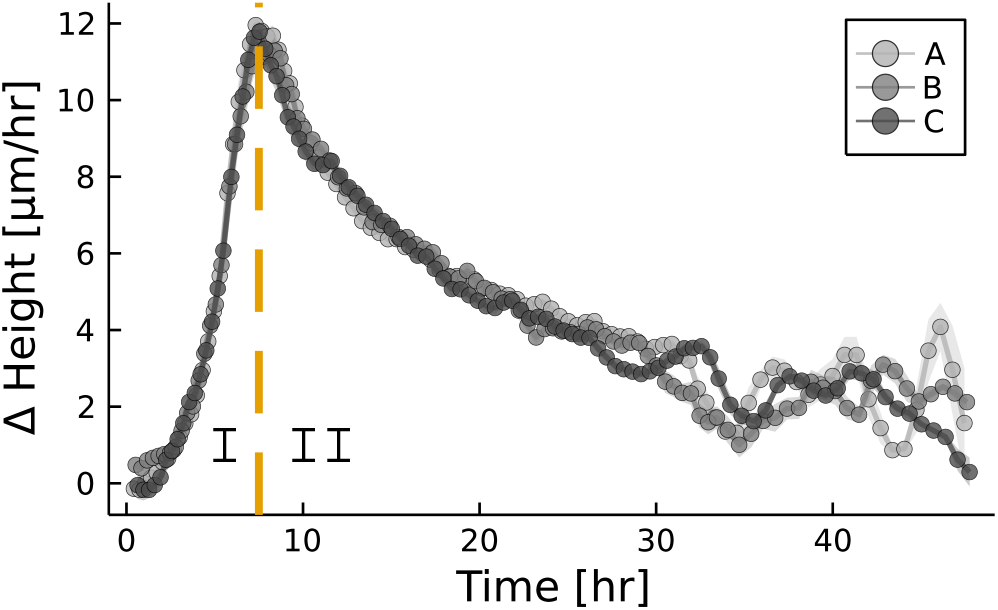
The growth regimes are not immediately evident when plotting their the growth over time. Each color represents a replicate of *A. veronii*, Measurements are the same as in the main figure.

**Figure S4.**
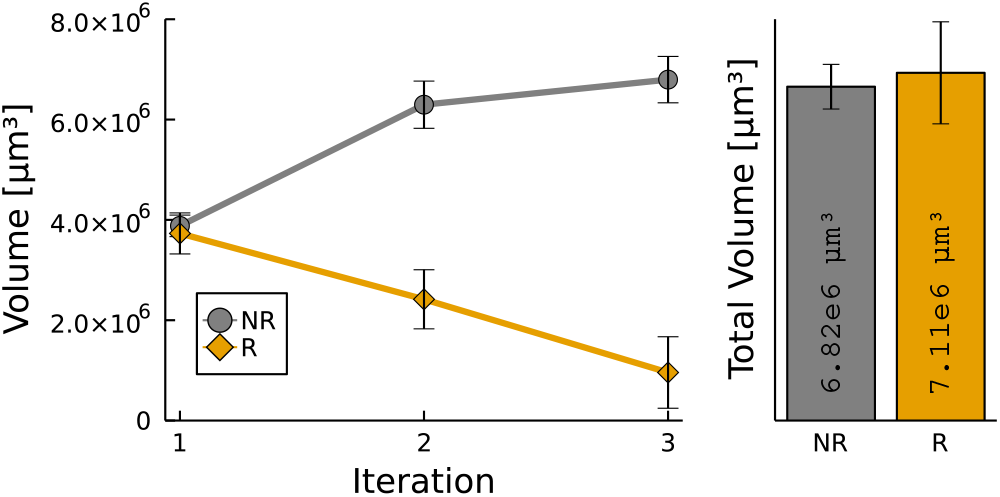
Growth of microbial colonies in a LB-Agar column, with (R) and without (NR) replacement every two days. (D) Total amount of growth supported by an LB-Agar colony over a period of 6 days.

**Figure S5.**
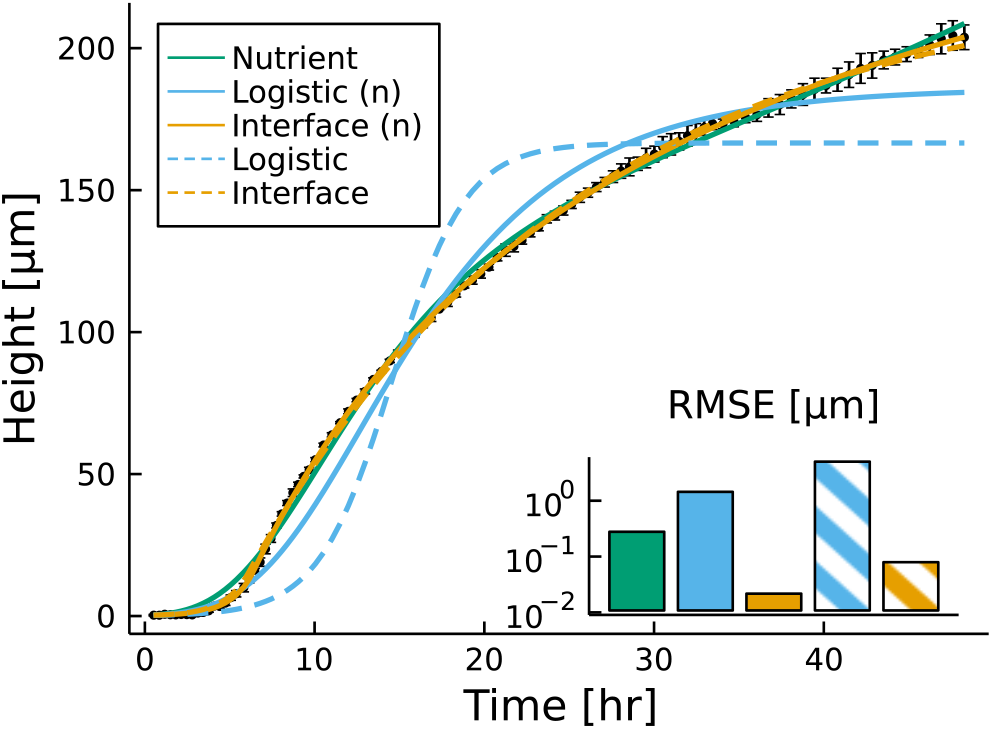
Best fits are shown for the different models while using a free optimization. For the nutrient model the parameters are: [3.3e7, −1.3e-1, 2.7e7, 5.2e6]. Logistic (n): [4.4e12, 5.2e2, 4.5e12, 5.9e11]. Interface (n): [2.0e0, 1.1e-1,1.6e1,1.8e0,-1.4e-1]. Logistic:[4.6e0, 1.6e2]. Interface:[7.5e-1,5e-2,1.5e1]. The parameter lists are ordered as in Table 2.

**Figure S6.**
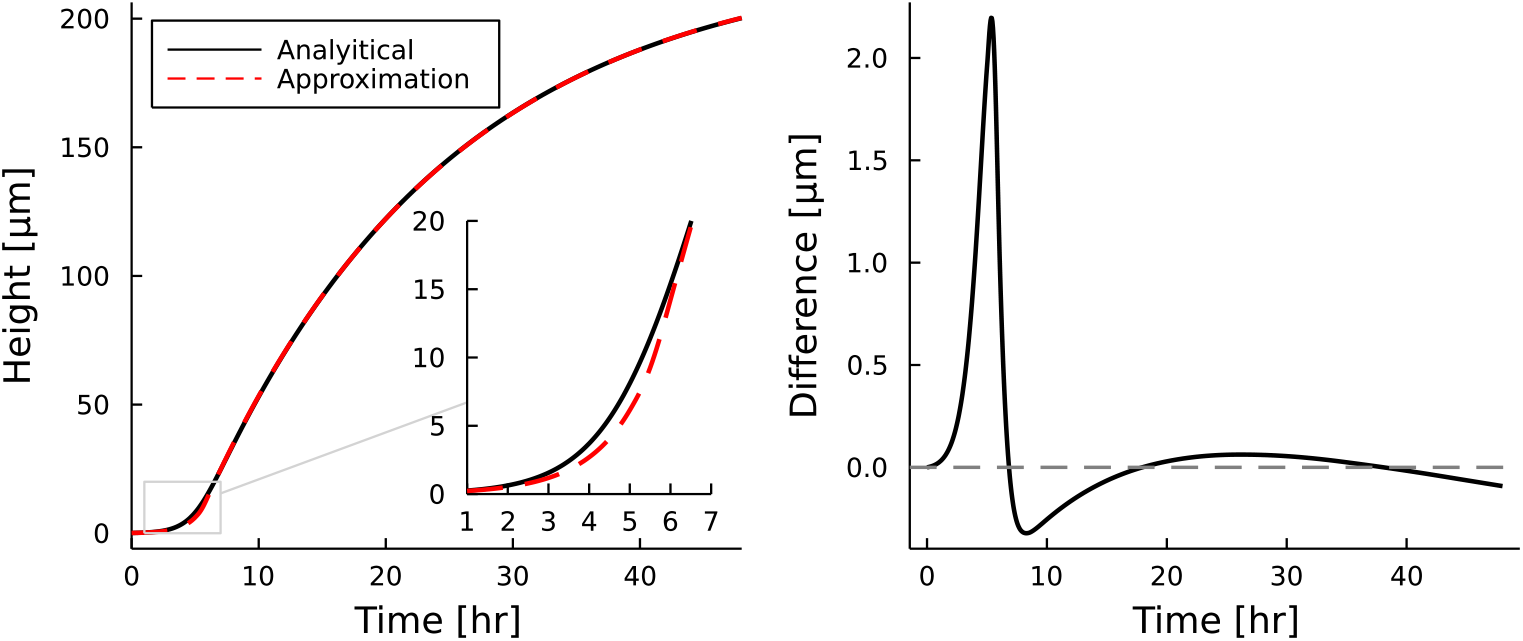
Best fits values are shown for the *A. veronii* timelapse in the main figure panel A. Parameters for the analytical expression are: 0.55, 0.05, 20.96, for the approximation: 0.87, 0.05, 13.20, for *α, β, L* respectively. The only significant discrepancies occur around the inflection point, but deviations do not exceed 2*µm* in magnitude. On the right, we see the difference between these models by plotting the analytical expression minus the approximation.

**Figure S7.**
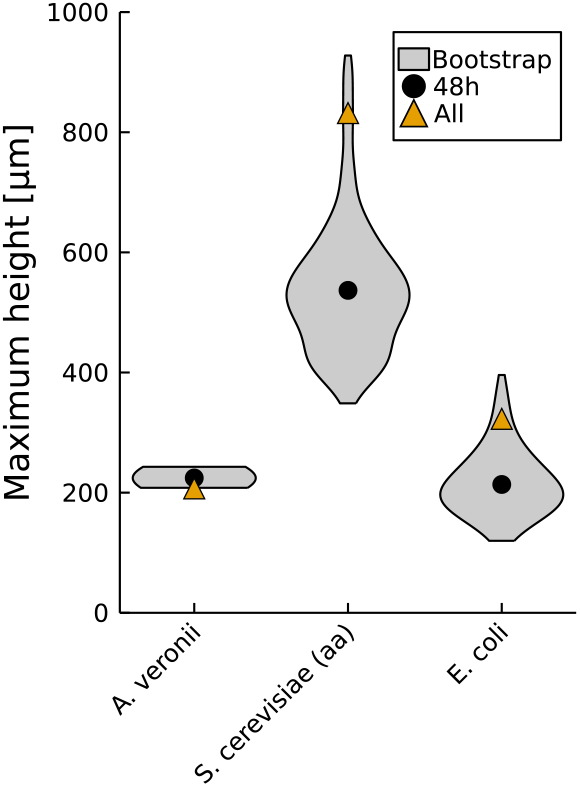
Interface model *h*_max_ distributions after N=1000 bootstrapped fits are shown. Black circles represent the best fit for the initial 48 hour time period, and orange triangles represent the best fit including the measurements up to 14 days.

**Figure S8.**
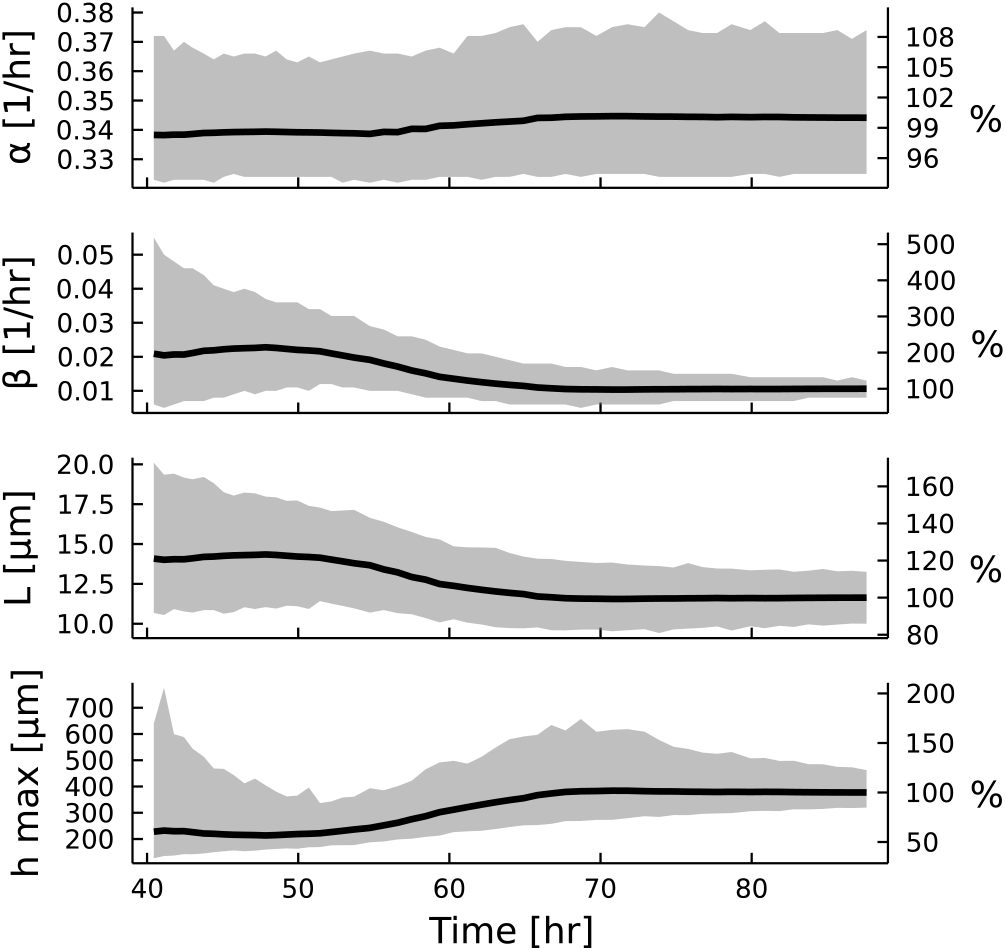
Best fit, and 90% confidence interval for the the three interface model parameters: *α, β, L*, and the steady state maximum height *h*_max_ are shown. On the left y-axis the actual values are reported, while on the right y-axis reports the value relative to the final 88 hour best fit value.

**Figure S9.**
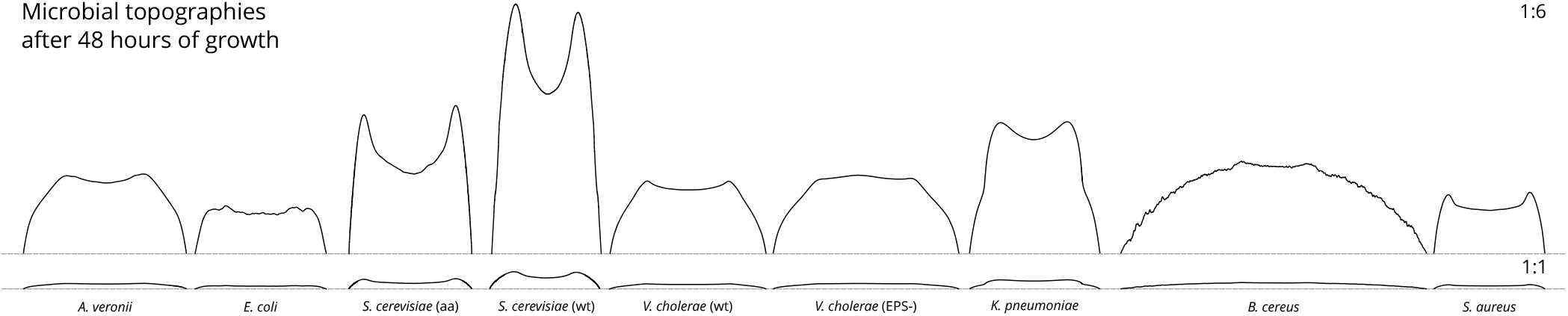
Profiles of the measured microbes after 48 hours of growth. Pictured are two different aspect ratios for the x and y axis for better visualization of the surface structures.

**Figure S10.**
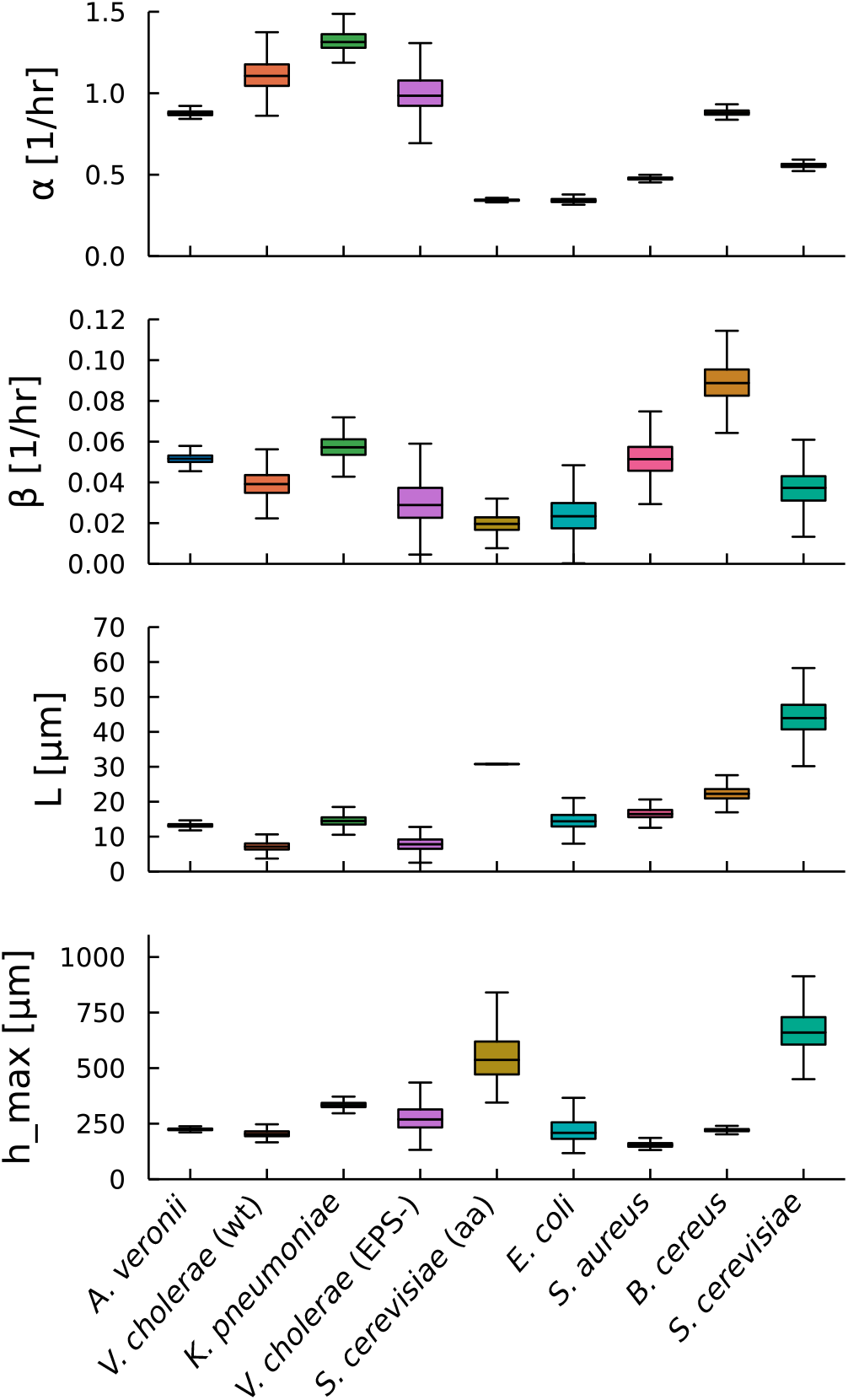
Fit parameters for the interface model across different bacteria. This panel demonstrate the range of possible parameter combinations, and similarities between strains.

**Figure S11.**
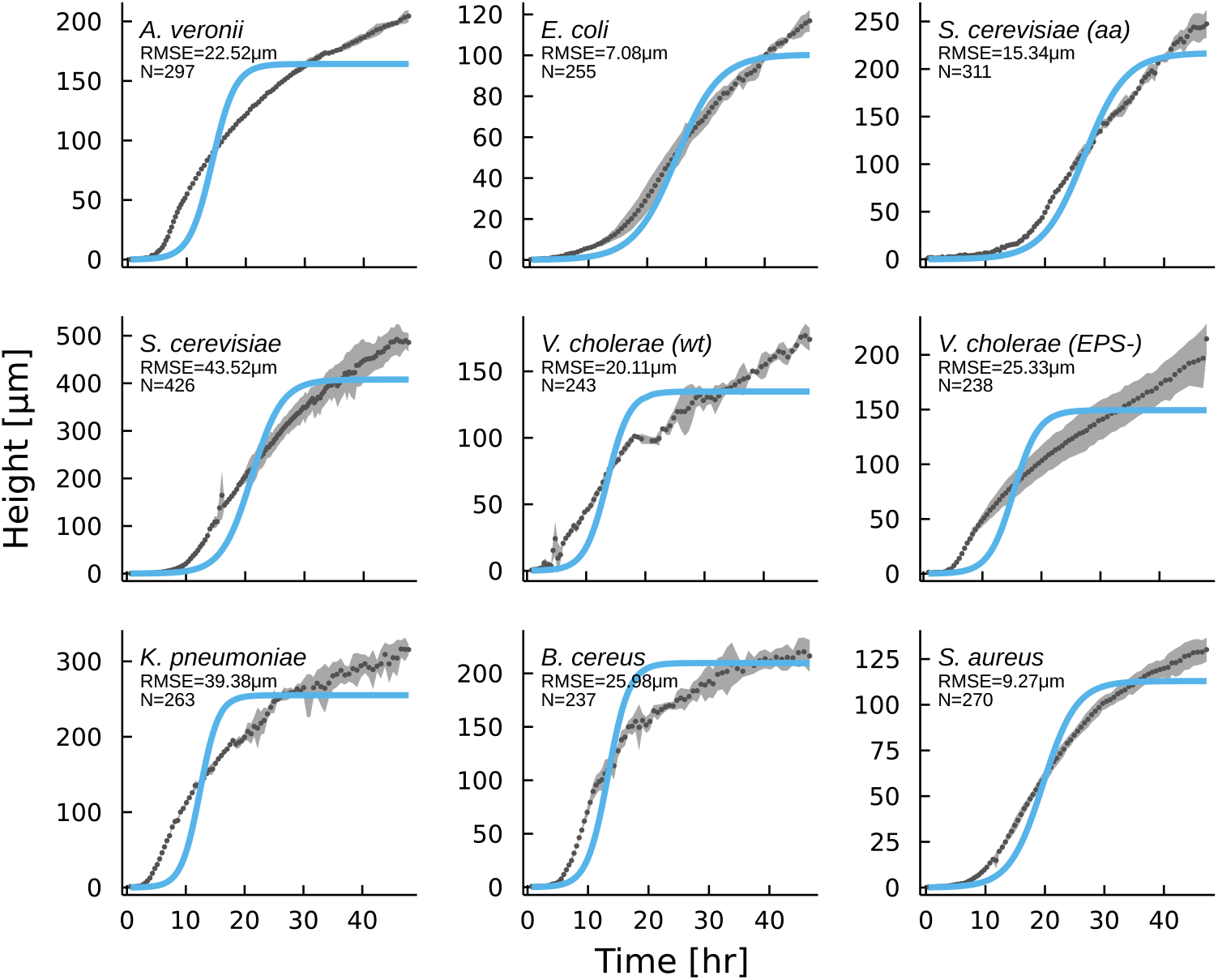
A simple logistic model fails to describe vertical height dynamics, across the sampled cohort of microbial colonies.

**Figure S12.**
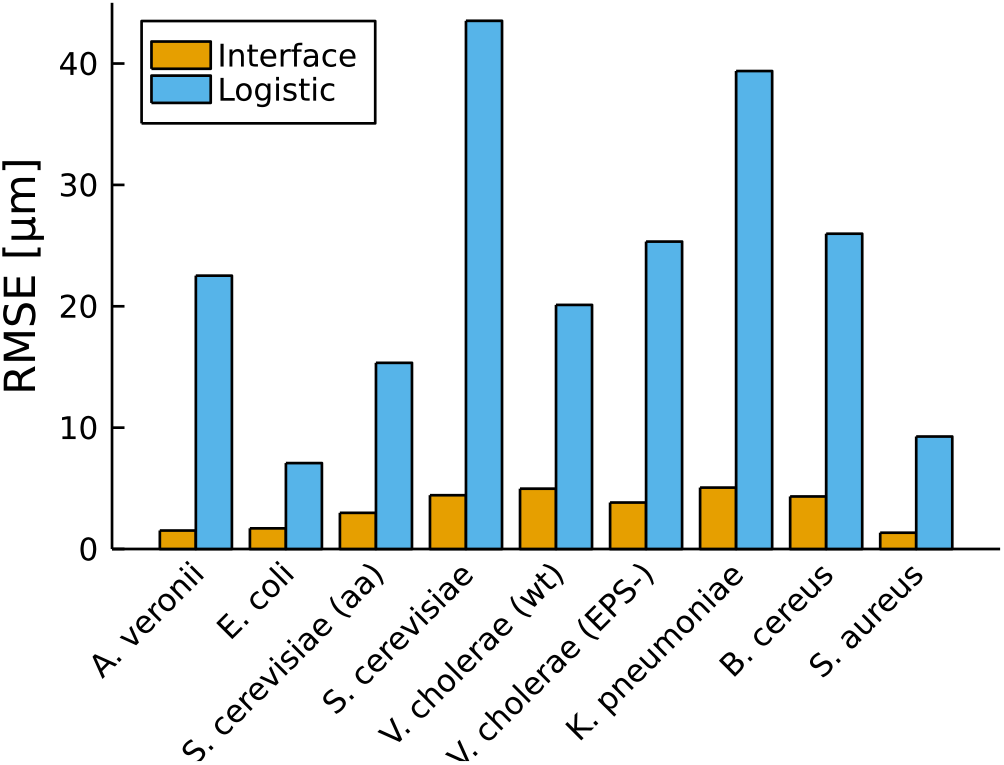
The interface model outperforms the logistic model in all sampled colonies.

## Notes

https://github.com/pbravoc/vertical_growth_biofilms

### Competing Interest Statement

The authors have declared no competing interest.

### Summary of Updates

The text was updated and expanded.

